# Cell Modeling and Rescue of a Novel Non-coding Genetic Cause of Glycogen Storage Disease IX

**DOI:** 10.1101/2025.05.14.654043

**Authors:** Apoorva K. Iyengar, Xue Zou, Jian Dai, Rhodricia A. Francis, Alexias Safi, Karynne Patterson, Rebecca L. Koch, Shannon Clarke, M. Makenzie Beaman, Jessica X. Chong, Michael J. Bamshad, William H. Majoros, R. Catherine Rehder, Deeksha S. Bali, Andrew S. Allen, Gregory E. Crawford, Priya S. Kishnani, Timothy E. Reddy

**Affiliations:** Department of Biostatistics and Bioinformatics, Duke University School of Medicine, Durham, NC 27710, USA; Biochemical Genetics Laboratory, Duke University Health System, Durham, NC 27710, USA; Division of Medical Genetics, Department of Pediatrics, Duke University School of Medicine, Durham, NC 27710, USA; Department of Genome Sciences, University of Washington, Seattle, WA 98195, USA; Department of Pathology, Duke University School of Medicine, Durham, NC 27710, USA; Department of Pediatrics, University of Washington, Seattle, WA 98195, USA; Brotman Baty Institute, Seattle, WA 98195, USA; Seattle Children’s Hospital, Seattle, WA 98105, USA; Center for Statistical Genetics and Genomics, Duke University, Durham, NC 27710, USA

**Author notes:** denotes co-corresponding authors: Timothy E. Reddy; 101 Science Drive, 2347 FCIEMAS Building, Durham, NC 27710, Gregory E. Crawford; 101 Science Drive, 2111 FCIEMAS Building, Durham, NC 27710, Priya S. Kishnani; 905 LaSalle Street, 4010 Genome Sciences Research Building 1, Durham, NC 27710.

## Abstract

Delayed diagnosis of Mendelian disease substantially prevents early therapeutic intervention that could improve symptoms and prognosis. One major contributing challenge is the functional interpretation of non-coding variants that cause disease by altering splicing and/or gene expression. We identified two siblings with glycogen storage disease (GSD) type IX γ2, both of whom had a classic clinical presentation, enzyme deficiency, and a known pathogenic splice acceptor variant on one allele of *PHKG2*. Despite the autosomal recessive nature of the disease, no variant on the second allele was identified by gene panel sequencing. To identify a potential missing second pathogenic variant, we completed whole genome sequencing (WGS) and detected putative deep intronic splicing variant in PHKG2 in both siblings. We confirmed the functional splicing effects of this variant using short-read and long-read RNA-seq on patient blood and a HEK293T cell model in which we installed the variant using CRISPR editing. Using the cell model, we demonstrated multiple biochemical and cellular impacts that are consistent with GSD IX γ2, and a reversal of aberrant splicing using antisense splice-switching oligonucleotides. In doing so, we demonstrate a novel and robust pathway for detecting, validating, and reversing the impacts of novel non-coding causes of rare disease.

## Introduction

Determining the genetic variants that cause Mendelian disease is a crucial step in accurate diagnosis and consequently in patient care. A prolonged diagnostic odyssey is common and has lasting effects on the physical, psychological, and financial wellbeing of patients and their families (1,2). Understanding the genetic and molecular mechanisms underlying a patient’s disease can inform prognosis, improve disease management, and may be required for insurance reimbursement and eligibility for clinical trials (3). Identifying novel causes of rare disease can also reveal new therapeutic targets for both rare and common disease. For those reasons, identifying additional genetic causes of rare disease is a profound opportunity for advancing precision medicine and improving healthcare (4–7).

Glycogen storage diseases (GSDs) (incidence: 1:20,000-43,000 live births) provide an instrumental example of that diagnostic odyssey. GSDs are a group of mostly autosomal recessive disorders caused by genes involving glycogen synthesis and breakdown, typically in liver and muscle cells (8,9). These inborn errors of carbohydrate metabolism have high genetic and phenotypic heterogeneity with symptoms ranging from exercise intolerance to liver failure; however, most are progressive and in severe cases can cause metabolic crisis and irreversible organ damage if left untreated (10). Identifying genetic variants that cause GSDs can lead to accurate diagnosis prior to the onset of severe symptoms, allowing early nutrition and other medical interventions that can delay or prevent major organ damage. In contrast, delays in diagnosis can lead to much worse outcomes in the short term and over a lifetime (11–13).

One of the major challenges in identifying novel causes of rare diseases, including GSDs, is the identification of variants that disrupt mRNA splicing, which are thought to be involved in at least 10% of Mendelian disease cases (14–18). That challenge persists for several reasons. On one hand, whole-exome sequencing (WES) – commonly used for diagnosing genetic disease – typically only identifies coding variants and non-coding variants at known splice sites that immediately flank exon boundaries. That inability of WES to identify a wide range of non-coding genetic variants may explain why the yield of genetic diagnosis from WES is <30% when monogenic disease is suspected (19–22). On the other hand, whole-genome sequencing (WGS) allows clinicians to observe nearly all non-coding variants. However, computationally predicting the impacts of variants outside of known splice sites, in particular the inclusion of deleterious pseudoexons in a transcript, remains a major challenge (23–27). For that reason, computational pseudoexon prediction alone is currently not considered sufficient to classify a deep intronic variant as pathogenic or benign (25,28).

Identifying pseudoexons that cause rare disease also has the potential to inform novel therapeutics. For example, the aberrant splice sites can be targeted for exclusion from the mRNA using splice-switching antisense oligonucleotides (SSOs) (29–31). Such SSOs have previously been shown to correct aberrant splicing in neurological diseases including spinal muscular atrophy and Batten disease (Neuronal Ceroid Lipofuscinosis). SSOs function through steric blocking of splicing factors from unwanted splice sites (32,33). GSDs are an excellent potential application of SSO technology because the liver is one of the few organs to which it is possible to deliver SSOs at sufficient concentrations; and because N-acetylgalactosamine (GalNAc) oligonucleotide modifications further enhance targeting to hepatocytes (34–36).

Here we describe a case study in which we discover a novel deep intronic variant that results in a pseudoexon in *PHKG2,* causing GSD IX γ2 symptoms in two siblings. We validated this variant by CRISPR editing a model cell line and demonstrated a candidate SSO that rescues wild-type *PHKG2* splicing. This study demonstrates a potential pipeline for improving rare disease diagnosis by using commonly available biosamples, cell models, and experimental technologies. In particular, we find that short-read and long-read RNA-seq are especially valuable for identifying splicing-related causes of rare disease; and that the creation of personalized cell culture models using genome editing enables a wide range of molecular and cellular assessments without requiring invasive collection of patient tissue. While many challenges remain in developing personalized therapy for rare diseases, these findings are a step towards making that process more routine.

## Results

### Clinical presentation of two siblings with glycogen storage disease IX γ2

We diagnosed two siblings with GSD IX γ2, which is an autosomal recessive disease caused by pathogenic variants on both alleles of *PHKG2*. This gene encodes the gamma subunit of hepatic phosphorylase kinase (PhK). This is a critical enzyme involved in glycogenolysis, responsible for activating glycogen phosphorylase and stimulating the release of glucose −1-phosphate in the liver, thus playing a crucial role in the regulation of blood glucose levels. Because it involves the catalytic gamma subunit, GSD IX γ2 is considered to be the most severe form of GSD IX, resulting in progressive liver injury including fibrosis or cirrhosis in >95% of patients, putting them at increased risk for requiring a liver transplant (37–39).

Sibling A’s parents are non-consanguineous and previously had two unaffected children. He presented at 9 months of age with delayed growth, hepatomegaly, hypoglycemia, and elevated liver ALT/AST and urinary glucose tetrasaccharides (Glc4). Despite careful dietary adherence, including uncooked cornstarch supplementation, high protein, and limiting fasting intervals to <4-6 hours including nights, he required a gastrostomy tube to maintain euglycemia. A liver biopsy revealed elevated glycogen content suggestive of a GSD, prompting an enzymology panel. PhK activity in the liver biopsy was 0.01 umol/min/g (reference range 0.14±0.11), consistent with GSD IX. Genetic testing via GSD IX gene panel sequencing found one known variant at *PHKG2* c.96-11G>A (40). This variant disrupts the splice acceptor of exon 3, resulting in a predicted deleterious SerSerCys insertion. That variant has been described as pathogenic or likely pathogenic (Variation ID: 1066974, 2 submitters, 1-2 star review status). At the beginning of this study, a second pathogenic, likely pathogenic, or variant of uncertain significance had not been detected.

Subsequent to diagnosis, sibling A’s liver symptoms continued to progress. He had progressive liver enlargement and increased echogenicity observed via ultrasound imaging and MRI (Figure 1A). Despite continuous glucose monitoring (Dexcom 6) confirming maintenance of euglycemia (Figure 1B), liver disease has continued to progress. Serum AST, ALT, and urine glucose tetrasaccharide (Glc4) – a biomarker of glycogen accumulation – were all initially elevated and have decreased over time (Figure 1C,D), coinciding with worsening features on liver imaging and biopsies showing fibrosis and cirrhosis (41,42).

**Figure 1.**
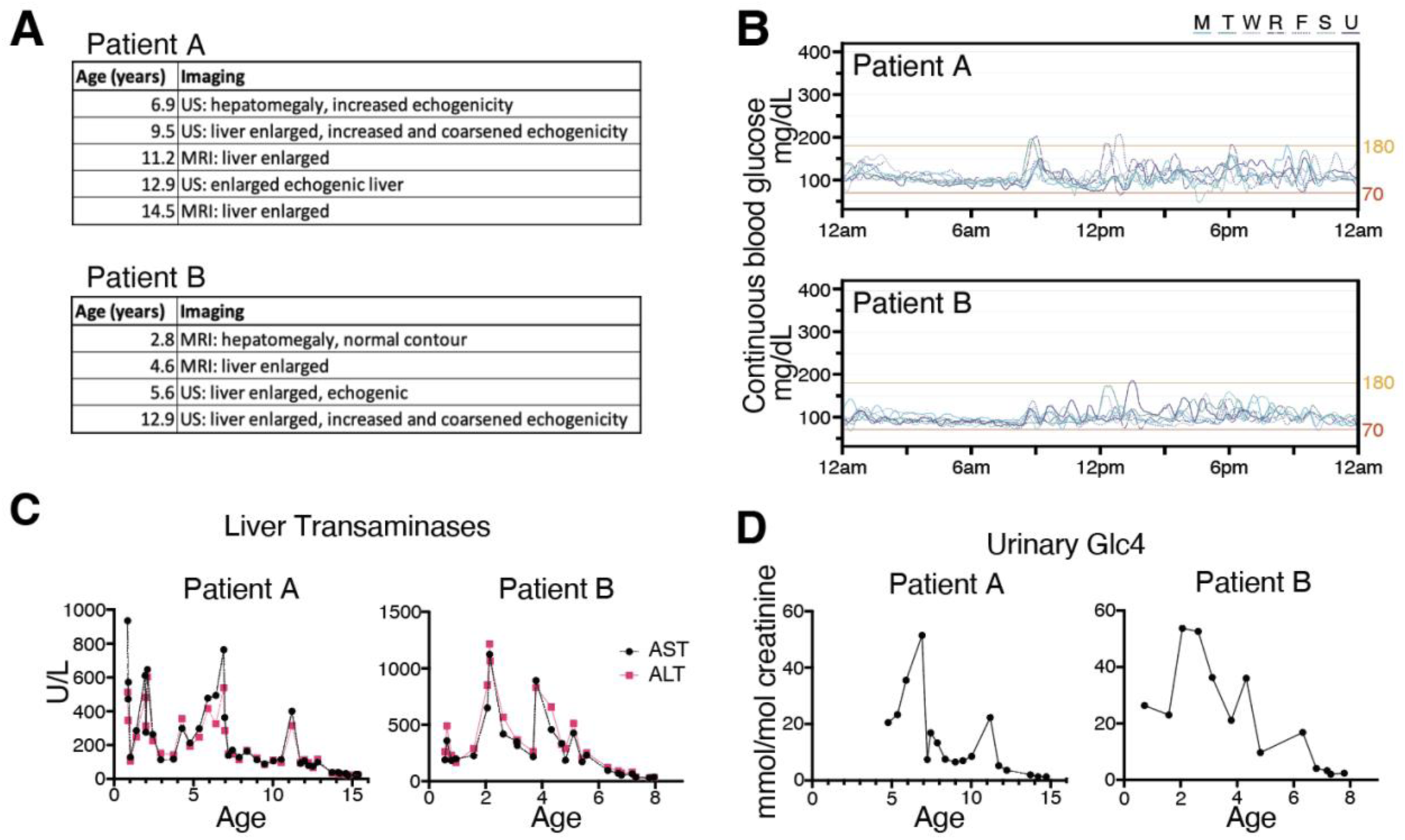
Management and progression of GSD IX γ2 in two siblings. (A) Liver imaging via MRI (Magnetic resonance imaging) and ultrasound (US) indicate hepatomegaly, and progressive fibrosis. (B) Continuous blood glucose monitoring over a representative week indicates that blood glucose is typically within normal limits (70-180 mg/dL) for both siblings. (C) Levels of aspartate and alanine aminotransferase (AST and ALT, respectively) have decreased over time, which is typical of progressive liver disease. (D) Urinary Glc4, a biomarker of glycogen accumulation, has decreased to normal levels.

Sibling B, born eight years later, was diagnosed with GSD IX γ2 at one month of age after genetic testing due to Sibling A’s clinical history. He also had delayed growth, hepatomegaly, and hypoglycemia requiring a gastrostomy tube. His liver biopsy showed low PhK activity at 0.02 umol/min/g (reference range 0.14±0.11). Like his brother, he carried the same likely pathogenic splice acceptor variant *PHKG2* c.96-11G>A, and no second variant was found.

Like Sibling A, Sibling B has also demonstrated progressive liver disease after diagnosis. Liver biopsies revealed enlarged hepatocytes, foci of chronic inflammation, and fibrosis beginning at age 2, with increased bridging and nodularity in subsequent biopsies. His serum and urine biomarkers were initially elevated and have decreased with time, and ultrasound imaging at ages 2, 4, and 5 years showed progressive hepatomegaly and echogenicity.

### Identification of a novel pseudoexon impacting PHKG2

GSD IX gene panel testing (*PHKA2*, *PHKB*, *PHKG2*) for both siblings only revealed a single likely pathogenic variant (c.96-11G>A) on one allele of *PHKG2* despite the autosomal recessive nature of GSD IX γ2. Based on the clinical phenotype and the low PhK enzyme activity, we hypothesized the siblings were compound heterozygous with a second unidentified variant in *PHKG2*. Because panel testing focused on exons and splice sites for those known exons, we also hypothesized that the missing variant was non-coding. Using whole genome sequencing (WGS), we detected one novel variant [NM_000294.3(PHKG2):c.556+1069T>G] in both patients that creates a cryptic splice donor site 1,069 bp downstream of exon 6. Using SpliceAI, we predicted with moderate to high confidence that this variant would create a 76 bp pseudoexon insertion in intron 6 (Donor Gain Δ=0.91, Acceptor Gain Δ=0.78, Figure S1) (43). If confirmed, that insertion would cause an out of frame, non-functional gene product via creation of a premature termination codon in the kinase domain of PHKG2.

We verified the predicted effects of both c.96-11G>A and c.556+1069T>G via RNA-seq on buffy coats obtained from both siblings. For c.96-11G>A, both siblings expressed a 9 bp extension to the *PHKG2* exon 3 splice acceptor, experimentally validating the three amino acid SerSerCys insertion predicted previously (Figure 2A, B) (44). Both siblings also expressed the 76 bp pseudoexon within intron 6 that we predicted for the c.556+1069T>G variant (Figure 2C). That pseudoexon was predicted to cause nonsense-mediated decay because it creates a premature stop codon that is more than 50-55 bp upstream of the 3’ exon junction complex (45). However, we did not observe significantly reduced *PHKG2* expression in either patient compared to the GTEx cohort of healthy individuals (Figure S3). We were unable to determine whether the allele containing c.556+1069T>G showed decreased expression due to the absence of heterozygous sites in the spliced mRNA and the distance between the two splicing events, which exceeded the read length.

**Figure 2.**
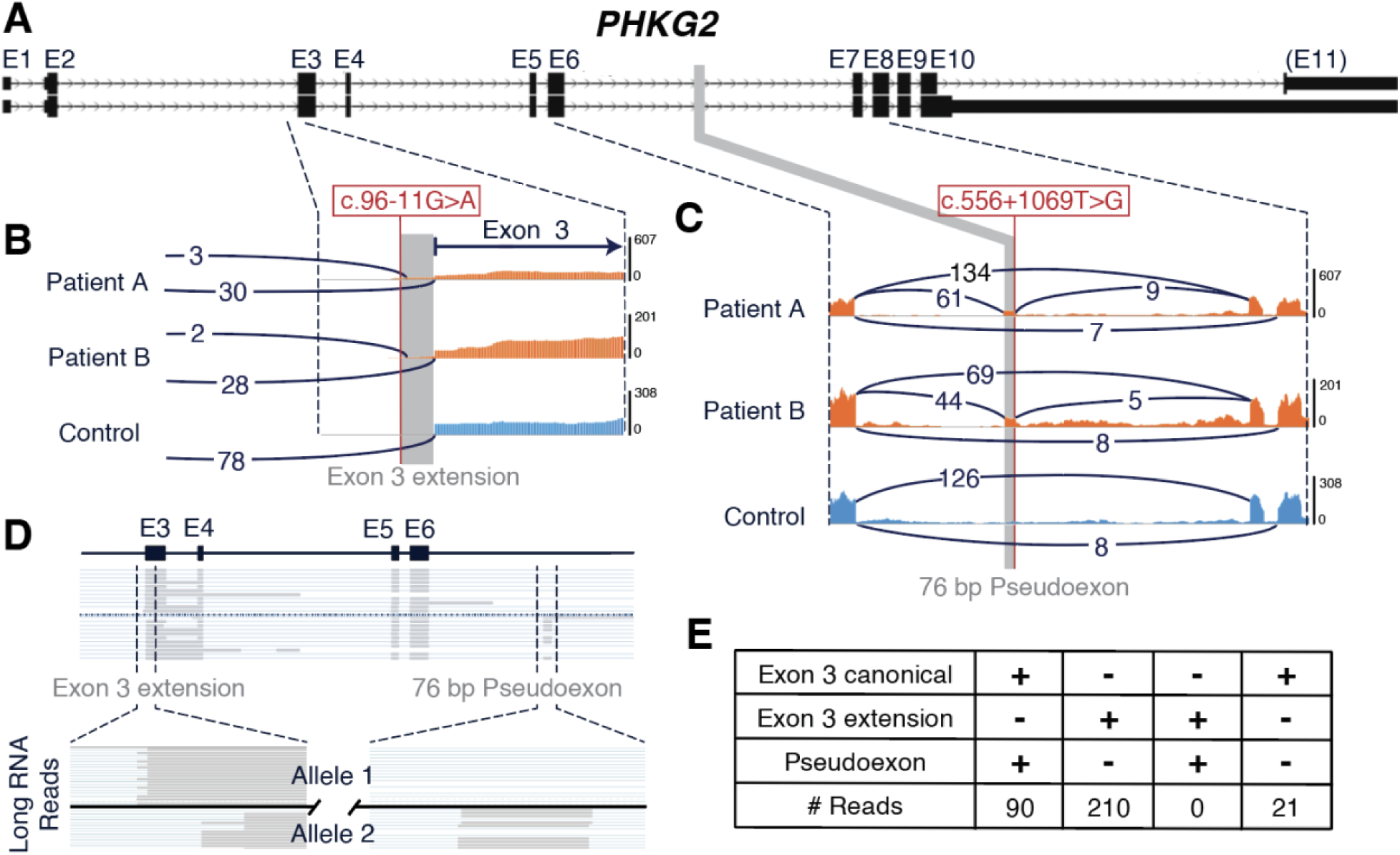
RNA sequencing of buffy coat from two siblings with GSD IX γ2. (A) Model of the *PHKG2* gene. (B) Short-read RNA-seq from buffy coat zoomed into the region (chr16:30,762,387-490, hg38) surrounding the originally identified c.96-11G>A variant. Both siblings show an alternate splice junction reflecting a 9 bp exon 3 extension in *PHKG2* near a splice site created by c.96-11G>A. This splice junction is not seen in a representative healthy control. (C) Short-read RNA-seq from buffy coat zoomed into the region (chr16:30,764,715-30,767,946, hg38) containing the newly identified deep intronic c.556+1069T>G variant. Both siblings express a novel 76 bp pseudoexon (gray box) with c.556+1069T>G at the cryptic splice donor. The pseudoexon is not detected in an unaffected control. (D) Targeted long-read RNA-seq show of Patient B. Reads containing the 9 bp exon extension are sorted above solid line labeled Allele 1. Reads containing the canonical exon 3 splice acceptor, some of which contain the 76 bp pseudoexon, are located on different transcripts shown below the solid line and labeled Allele 2. (E) Number of long-read RNA-seq transcripts containing either the exon 3 extension (due to c.96-11G>A) or the pseudoexon (due to c.556+1069T>G). There are no reads containing both the exon 3 extension and the pseudoexon, indicating that the causal variants are in *trans*. With 90 reads mapping to the pseudoexon and 210 reads mapping to the exon 3 extension, the expression profile indicates allelic imbalance.

To confirm that c.96-11G>A and c.556+1069T>G are compound heterozygous variants, we tracked the origins of the alleles in the parents. To do so, we used RNA-seq instead of DNA sequencing to directly observe the consequences of each of the two *PHKG2* variants on different mRNA transcripts. The mother only expressed the exon 3 extension consistent with c.96-11G>A (Figure S2A), while the father only expressed the 76 bp pseudoexon consistent with c.556+1069T>G, indicating that the two variants are in *trans* in both patients (Figure S2B). We further investigated the expression effects of these variants using targeted long-read RNA-seq of Patient B (Figure 2D). In that analysis, 321 long-read RNA-seq reads spanned the sites of both c.96-11G>A and c.556+1069T>G. None of those reads contained both the exon 3 extension and the 76 bp pseudoexon, further confirming compound heterozygosity (Figure 2E). Overall *PHKG2* expression was within the normal range for both patients and parents (Figure S3). However, we identified imbalanced allelic expression that is consistent with nonsense-mediated decay as a result of the predicted premature termination codon caused by c.556+1069T>G (Figure 2D,E).

Of the 321 reads that span both aberrant splice sites, 210 (65%) contain the in-frame 9 bp exon 3 extension, 90 (28%) contain the out-of-frame 76 bp pseudoexon, and 21 (7%) are normal at both sites and encode the functional version of *PHKG2 (*Figure 2E). Therefore, although RNA-seq showed normal *PHKG2* expression compared to the GTEx cohort, the majority of transcripts from these patients lead to nonfunctional or no protein product. Furthermore, the low percentage of mRNA reads without aberrant splicing may explain the low but non-zero levels of PhK enzyme activity in the patients (Table 1).

### Patient-specific model of a novel pseudoexon causing GSD IX

To confirm that c.556+1069T>G is the cause of the 76 bp pseudoexon inclusion, we used CRISPR/Cas9 genome editing to create a model of the variant in HEK293T cells (Figure 3A). Biologically independent clones were generated that were either homozygous for the wild-type allele (T/T) or the patient allele (G/G) at this variant, allowing us to test the effects of the variant compared to an isogenic wild-type control. Genotypes were confirmed by allele-specific PCR and Sanger sequencing (Figure S4).

**Figure 3.**
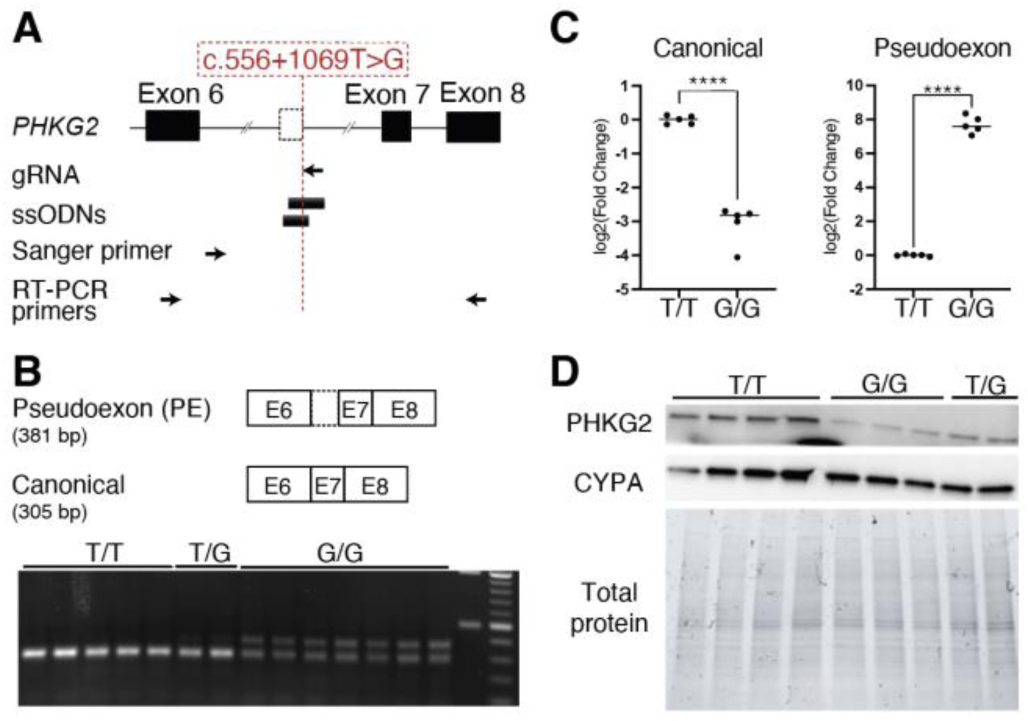
Creation of a patient-specific cell model of GSD IX γ2. (A) Schematic of gRNA and single stranded oligo donors (ssODNs) used to create *PHKG2* c.556+1069T>G edit and primers used to validate the cell line sequence and the presence of the pseudoexon. (B) RT-PCR of the region surrounding the 76 bp pseudoexon shows the addition of the larger 380 bp band, reflecting pseudoexon inclusion only in cell lines carrying the edited T>G variant (G/G alleles) relative to the wild-type cells (T/T alleles). (C) RT-qPCR quantification of the canonical isoform measuring expression of the Exon 6-7 junction shows an 89% decrease of the canonical PHKG2 transcript in cells containing c.556+1069T>G. Quantification of the Exon 6-pseudoexon junction shows significant expression only in c.556+1069T>G edited cells. (D) Western blot of PHKG2 shows decreased protein expression in c.556+1069T>G edited cells. (****p<0.0001)

RT-PCR across the site of this variant (Figure 3A) showed only the canonical 305 bp isoform band in wild-type (T/T) cells, while edited (G/G) cells expressed the canonical isoform in addition to a second 381 bp band corresponding to a 76 bp pseudoexon-containing isoform (Figure 3B). Sanger sequencing of the 380 bp band confirmed the 76 bp pseudoexon inclusion between exons 6 and 7 (Figure S5). We quantified the relative abundance of the isoforms using probe-specific RT-qPCR, with the qPCR probe spanning either the exon 6-7 boundary or the exon 6-pseudoexon boundary. This indicated an 87% decrease in canonical *PHKG2* expression (Figure 3C, p<0.0001) and 52% decrease in overall *PHKG2* expression (Figure S6, p<0.001) in the edited (G/G) cells compared to wild-type (T/T), providing evidence that c.556+1069T>G and its resulting pseudoexon causes nonsense-mediated decay. This decrease in PHKG2 persists at the protein level as measured by a western blot (Figure 3D).

To our knowledge, this is the first cellular model of GSD IX generated via genome editing of non-patient cells. That approach has the key advantage that it provides isogenic controls for the individual variant of interest. We therefore sought to characterize disease-relevant phenotypes to determine whether it may be an appropriate model to gain insight into GSD IX etiology and for testing potential therapeutics. Biochemical testing of PhK enzyme activity, similar to that used in erythrocytes or liver biopsies to diagnose GSD IX, showed a 90% reduction in PhK activity in HEK293T cells containing c.556+1069T>G (p<0.0001, Figure 4A). This is concordant with the reduced level of PhK activity in the GSD IX patients.

**Figure 4.**
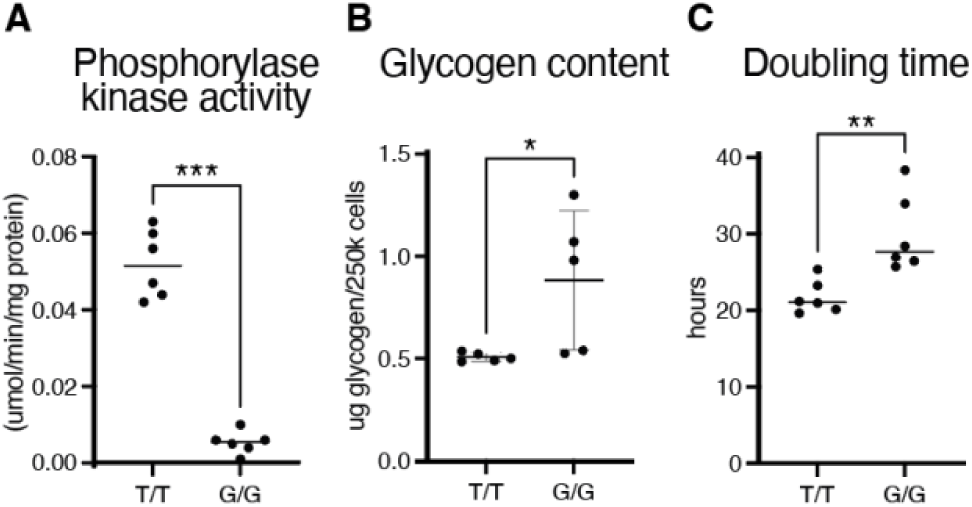
Characterization of GSD IX γ2 HEK293T cell line containing the c.556+1069T>G variant. (A) Clinical testing for phosphorylase kinase enzyme activity of c.556+1069T>G edited cells (G/G alleles) display a 10-fold decrease in phosphorylase kinase activity *in vitro* compared to wild-type cells (T/T alleles), a similar level of change detected in cases compared to controls in human blood cells and liver biopsies (Table 1). (B) In cells grown under low glucose conditions to induce glycogenolysis, glycogen content is higher in c.556+1069T>G edited cells compared to wild-type. (C) Cell doubling time is significantly higher and cell count significantly lower in c.556+1069T>G edited cells than wild-type for 48 hours under low-glucose conditions. (*p<0.05, **p<0.01, ***p<0.001)

Because PhK is a critical component of the glycogenolysis pathway, we hypothesized that the HEK293T cells with c.556+1069T>G variant would have reduced ability to metabolize stored glycogen. To create an environment that would promote glycogenolysis, cells were cultured in media with reduced glucose (0.5 g/L) and pyruvate (55 mg/L), the product of glycolysis. After 24 hours, glycogen content was higher in c.556+1069T>G HEK293T cells compared to wild-type cells (p<0.05, Figure 4B). After allowing the cells to grow under the same conditions for 48 hours, the calculated doubling time of the c.556+1069T>G cells was lower than wild-type cells (Figure 4C). This effect could not be attributed to differences in the rate of apoptosis, indicating that blunted cellular metabolism in c.556+1069T>G cells may slow energy-intensive functions including cell replication (Figure S7).

### Classification of *PHKG2* c.556+1069T>G

Numerous lines of evidence from population data, functional experiments, allelic effects, and patient phenotypes indicate that this variant is pathogenic as determined by ACMG variant classification guidelines, including updated recommendations for non-coding variants (25,28). The variant is absent from gnomADv4 with >450,000 alleles covering the region; thus, we apply PM2 at the supporting level (46). As confirmed by splicing analysis of parental RNA-seq and long-read RNA-seq of Patient B, this variant is in *trans* with the previously known pathogenic variant c.96-11G>A, allowing us to apply PM3 with a moderate level of evidence. Biochemical evidence of low PhK activity in liver biopsies combined with a clinical presentation of hypoglycemia, hepatomegaly, and progressive liver fibrosis is highly specific to GSD IX, and the level of severity of the patients’ disease is concordant with variants in the γ2 subunit encoded by *PHKG2* (37). Thus, we can apply PP4 at the moderate level. This variant is predicted by SpliceAI to create a 76 bp pseudoexon, and this prediction is validated by RNA-seq in multiple patients without evidence of compensatory splicing elsewhere in the gene. In addition, the variant causes an 87% decrease in splicing of functional *PHKG2* corresponding to a 90% decrease in PhK enzyme activity in a gene edited cell model. With this evidence of a near-complete loss of normal splicing, we can apply PS3 at the strong level. Together, these lines of evidence (one strong, two moderate, and one supporting) indicate that c.556+1069T>G should be classified as pathogenic.

### Rescue of canonical splicing via splice-switching antisense oligonucleotides (SSOs)

Splicing defects can be targeted by a class of therapeutics known as splice-switching oligonucleotides (SSOs). These antisense oligonucleotides base-pair with a pre-mRNA molecule to inhibit RNA-RNA base pairing events or splicing factor binding events that are critical for the aberrant splicing event to occur (47). Successfully blocking those interactions prevents the aberrant splicing and restores the normal mRNA sequence, reading frame, and amino acid sequence of the protein (48,49).

We designed three SSOs to inhibit pseudoexon inclusion by blocking the splice donor site (SSO1), a predicted exonic splicing enhancer as defined by a cluster of SR protein binding motifs (SSO2), and the splice acceptor site containing c.556+1069T>G (SSO3)(Figure 5A). Each SSO was 24-25 nucleotides with their sequences optimized to improve cellular uptake, avoid off-target effects, and avoid self-complementarity and G-quartet structures that reduce binding potential of the oligonucleotide (50). All SSOs had phosphorothioate backbones to increase stability and a 2’ O-Methyl (2’ O-Me) modification to resist nuclease degradation (51).

**Figure 5.**
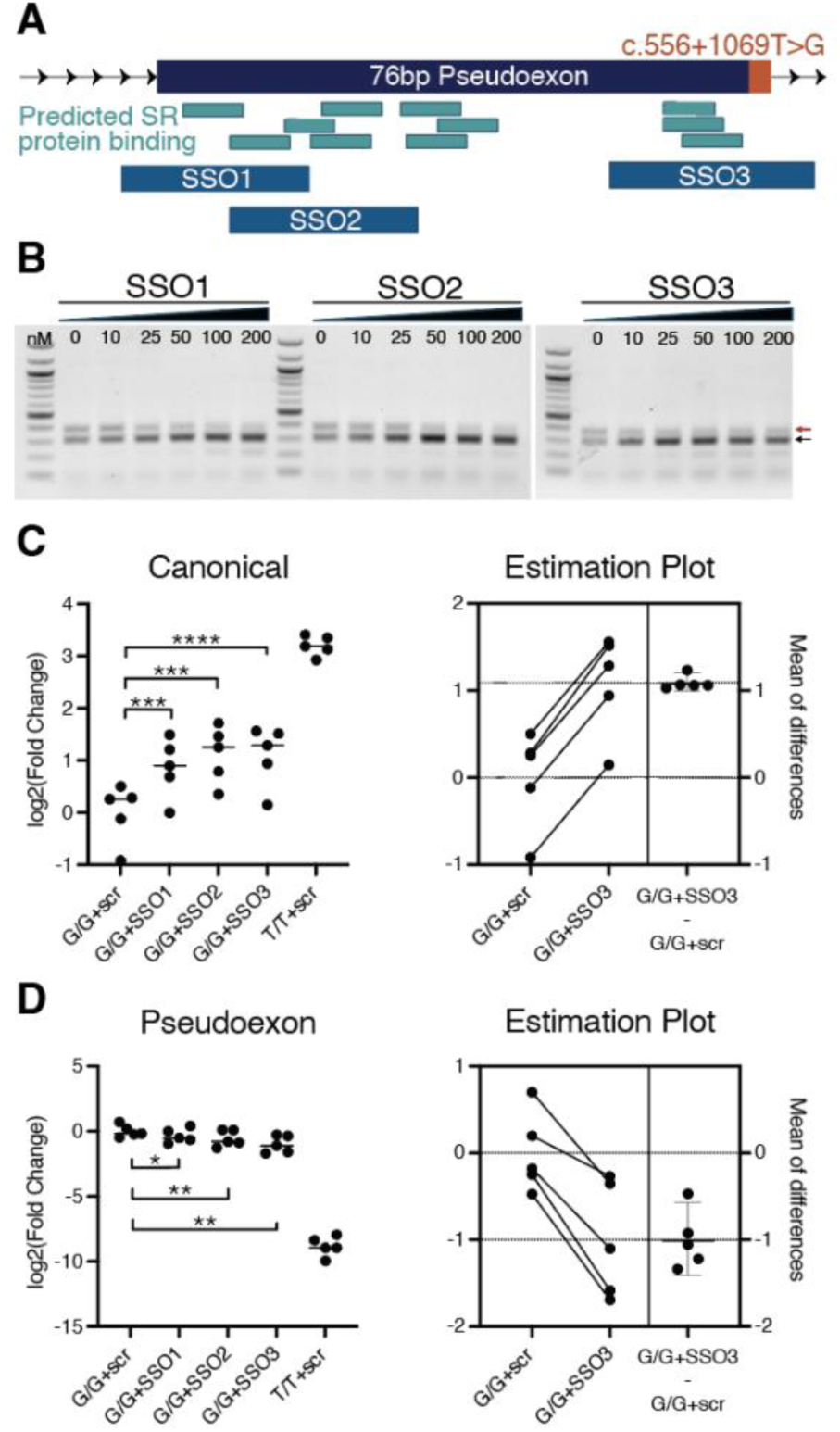
Design and rescue of canonical *PHKG2* expression via splice-switching oligonucleotides. (A) Three SSOs were designed at the 76 bp pseudoexon at the 5’ splice acceptor (SSO1), the 3’ splice donor (SSO3), and at a predicted exonic splice enhancer as defined by the presence of SR binding protein motifs (SSO2). (B) Each SSO was transfected into a representative c.556+1069T>G edited cell line. RT-PCR of the region surrounding the pseudoexon showed a dose-dependent rescue of the canonical isoform (black arrow) of PHKG2 and a corresponding decrease in the pseudoexon isoform (red arrow). All lanes were run on the same gel but were noncontiguous. (C) All 3 SSOs were transfected into 5 biologically identical GSD IX cell lines. RT-qPCR shows a significant increase in canonical isoform expression for all SSOs relative to a scrambled control SSO. SSO3 shows the largest rescue of the canonical isoform and decrease in pseudoexon isoform. The estimation plot indicates that the increase is consistent across all biological replicates. (D) For the same cell lines in (C) but quantifying the pseudoexon presence. The estimation plot indicates that pseudoexon inclusion decreased in each cell line. (*p<0.05, **p<0.01, ***p<0.001, ****p<0.0001)

The three SSOs were initially tested in a single homozygous c.556+1069T>G HEK293T clonal cell line to determine whether we could induce splice-switching at the pseudoexon locus. Each SSO was transfected in a dose-response curve (0-200 nM), and RT-PCR on RNA 24 hours post-transfection determined that all three SSOs rescued canonical isoform expression and decreased pseudoexon isoform expression in a dose-dependent manner (Figure 5B). The SSOs and matched scrambled controls were then transfected into five independently derived clonal c.556+1069T>G cell lines. All three SSOs showed significant rescue of canonical PHKG2 expression, with each biological replicate showing a rescue effect for all SSOs (Figure 5C,D). SSO3, targeting the cryptic splice site created by c.556+1069T>G, had the highest effect, improving expression of the functional isoform of *PHKG2* to 28% of that in wild-type cells. A version of SSO3 using the 2’-O-methoxyethyl (2’-MOE) modification, which is most commonly used in human therapeutic contexts due to higher tissue half-life and fewer pro-inflammatory effects, also significantly increased expression of the canonical *PHKG2* isoform (Figure S8) (32,33). Together, this indicates that SSOs partially restore normal PHKG2 splicing.

## Discussion

In this study, we identified and characterized a novel deep intronic pathogenic splice variant that was missed by panel sequencing in two GSD IX patients, by using a combination of WGS and RNA-seq. We validated the function of that variant by creating a patient-specific cell model of their disease using CRISPR genome editing. Finally, we characterized a potential therapeutic modality to reverse the splicing defect using SSOs. Together, these results demonstrate a pathway to discover, model, and revert novel non-coding pseudoexon inclusion events that cause rare disease.

The increased use of RNA-seq in recent years in patient samples for both rare and common disease has confirmed that pseudoexon retention is under-ascertained (26,29). Because pseudoexon detection is much easier from RNA, RNA-seq may be a high-yield 2nd line diagnostic tool when exome sequencing does not yield a genetic diagnosis. Long-read RNA-seq can be particularly useful for identifying pseudoexons and determining whether multiple pathogenic variants are on separate alleles. In our study, short-read RNA-seq with a two-pass alignment strategy was not able to consistently detect split reads at the novel splice junctions, with the pseudoexon splice acceptor being called six to eight times more frequently than the splice donor. It was also unable to resolve the exact length of the pseudoexon, making it unclear whether the splicing change led to a frameshift. In contrast, because long-read RNA-seq produced hundreds of reads spanning the majority of *PHKG2*, it was possible to assess functional effects of the pseudoexon inclusion, including 1) phasing the variants and splicing defects across the entire transcript, 2) identifying allele-specific expression, and 3) more accurately estimating the percentage of functional *PHKG2* mRNA expression. Therefore, in cases where only one or a few genes are of potential relevance, targeted long-read RNA-seq with high coverage of those genes can be of great use in identifying and classifying potential non-coding pathogenic variants.

After identifying variants of interest through sequencing and predicting their impact on protein function, a crucial next step in variant classification is verifying these predictions through functional assays (28,52). Classic minigene assays have verified the impact of numerous variants on deep intronic splicing (53,54). However, these assays are episomal and their scope is limited to one or two splice sites. Thus, they cannot functionally assess whether pseudoexon inclusion leads to nonsense-mediated decay or the degree to which it impacts protein function. With genome editing, we evaluated the splice variant both in its full genomic context and in an environment where disease-relevant phenotypes can be measured. Genome editing in non-patient cell models also has the advantage of being able to compare the gain of the candidate pathogenic variant to isogenic controls. An alternative and complementary strategy is to revert the candidate pathogenic variant in patient-derived and differentiated induced pluripotent stem cells (iPSCs) (55). Though there are clear benefits to both approaches, iPSCs have more technical challenges in that they are time-intensive, expensive, and can have low editing efficiency. In this study, we demonstrate that HEK293T cells may be a more scalable alternative as biochemical analysis of phosphorylase kinase activity in CRISPR edited cells showed that they express this key phenotype at a level concordant with what is expected in GSD IX patients (56). Since HEK293T cells are highly transfectable, editable, and divide quickly, they also have the potential benefit of possibly being used to assess dozens of variants at once.

Despite their limitations in modeling other GSD IX phenotypes, this study has also demonstrated that genome edited HEK293T cells can be used as a first-line tool to assess GSD IX therapeutics that target the genome or transcriptome, such as gene therapy or antisense oligonucleotide therapies. Multiple SSOs targeting the pseudoexon caused by c.556+1069T>G improved expression of the functional *PHKG2* isoform to 20-30% of that expressed by wild-type cells. Due to the rapid mitotic rate of HEK293T cells, evaluating the cellular impacts of transiently-delivered chemically modified SSOs was not possible, and the extent to which *PHKG2* expression must be rescued to bring glycogen phosphorylase activity above the pathogenic range is unclear. Therefore, further research is required to assess whether this level of rescue is adequate or the SSO sequences require further optimization.

GSD IX is primarily managed through strict dietary intervention, including high protein and complex carbohydrates, low simple carbohydrates, frequent small meals, and uncooked corn starch supplementation. Even highly compliant patients like those described in this study may experience severe hypoglycemia and substantial progressive liver fibrosis or cirrhosis. SSOs and other antisense therapies have substantially slowed disease progression in several diseases (30,57,58). Preliminary studies in GSD Ia and GSD II, in conjunction with the success of using GalNAc-conjugated oligonucleotides to target hepatocytes with systemic administration of the drug, indicate that SSOs are a promising strategy that could slow the progression of GSD IX (35,59). Although SSOs are not curative, they could substantially improve patient quality of life and in severe cases, delay the need for liver transplantation. This would be an excellent outcome due to the scarcity of donor organs, length of time before a donor organ becomes available, and the lifelong need for immunosuppression after organ transplant. As with many therapies, antisense oligonucleotides have the highest impact when deployed early in disease progression or even prior to the onset of symptoms, highlighting the need to develop strategies for detecting non-coding pathogenic variants (60,61).

## Methods

### Sex as a biological variable

One sex was involved in this study due to its focus on an extremely rare variant in two male patients. However, we performed our gene editing assays in a cell line derived from a female, and enzyme studies were consistent with those in the male patients. In addition, multiple lines of evidence for classification of c.556+1069T>G are relevant to both sexes. Thus, we expect that our findings with regards to the classification of are relevant for both sexes.

### Whole-genome sequencing and analysis

The genomic DNA library was constructed using the KAPA Hyper Prep kit per manufacturer protocol (KAPA Biosystems, #KR0961 v1.14). Massively parallel sequencing-by-synthesis with fluorescently labeled, reversibly terminating nucleotides was carried out on the NovaSeq sequencer. Reads were aligned the hg19 using BWA-MEM (Burrows-Wheeler Aligner; v0.7.15) with duplicate removal (Picard MarkDuplicates v2.6.0) and base quality recalibration (GATK BaseRecalibrator v3.7) (62). Variants were detected using the HaplotypeCaller tool from GATK v3.7 and annotated using the SeattleSeq Annotation Server (http://gvs.gs.washington.edu/ SeattleSeqAnnotation/) and manual curation in IGV. Candidate variant analysis was performed by annotating all variants with VEP and filtering on impact, inheritance, and allele frequencies using GEMINI (https://github.com/arq5x/gemini).

### Short-read RNA sequencing and analysis

RNA was prepared from frozen buffy coats through Trizol-chloroform extraction followed by isopropanol precipitation. Quantity was assessed on the Qubit fluorometer (ThermoFisher #Q32852) and RNA integrity was assessed on the 4200 TapeStation System (Agilent #5067). 1 ug of RNA was used to generate RNA-seq libraries using the TruSeq Stranded mRNA Library Prep kit. 50 bp paired end libraries were sequenced on a Novaseq 6000 S-prime flow cell. Fastq reads were mapped to the hg19 genome assembly using *STAR 2.7* in two-pass mode with WASP filtering in order to detect novel exon junctions (63).

Frozen whole blood PAXgene RNA samples were prepared using the PAXgene Blood RNA kit (Qiagen #762174) according to manufacturer’s instructions and quantity/quality were assessed as described above. Samples were globin depleted using GlobinClear-Human kit (ThermoFisher #AM1980) and rRNA depleted using Qiagen FastSelect HMR (Qiagen #334376). Libraries were prepared using the NEBNext Ultra II Directional RNA Library Prep Kit for Illumina (NEB #E7760), sequenced to a target depth of 300 million paired end 150 bp reads, and processed using the GTEx RNA-seq pipeline (https://github.com/broadinstitute/gtex-pipeline). To compare *PHKG2* expression to the non-globin depleted GTEx samples, *PHKG2* TPM was divided by *ACTB* or *B2M* TPM) for all samples. Splicing alterations were visualized in IGV.

### Long-read RNA sequencing and analysis

Full-isoform cDNA was prepared and PCR amplified (18 cycles) from 1 ug RNA using the SMARTer PCR cDNA Synthesis Kit (Takara #634925). 2.5 ug cDNA was probed with an IDT xGen Lockdown Pool of biotinylated oligonucleotides targeting GSD-related genes of interest (*ACADVL, AGL, ETFA, ETFB, ETFDH, GAA, GBA, PHKG2, SLC37A4*) and housekeeping genes (*ACTB, ATP5F1, PGK1).* 100 uL M-270 Streptavidin Dynabeads (ThermoFisher #65305) were washed according to manufacturer’s instructions and incubated with cDNA for 45 mins at 65°C to isolate biotinylated cDNA fragments. Captured cDNA was washed according to manufacturer’s instructions and amplified for 25 cycles using LA Taq DNA Polymerase Hot Start Version (Takara #RR042). The library was prepared using the SMRTbell Express Template Prep Kit 2.0 with barcoded adapters, omitting shearing and size selection. The resulting SMRTbell library was run for 24 hours on a Sequel I SMRT Cell. Circular consensus sequencing (CCS) reads were mapped to the hg19 genome assembly using minimap2 using default parameters of the PacBio minimap2 wrapper (pbmm2). Samples were assessed manually for novel splice junctions and for read numbers corresponding to allele-specific expression.

### HEK293T cell culture

HEK293T cells (ATCC) were maintained and gene editing was performed in high glucose media DMEM with 4.5 g/L glucose, 110 mg/L pyruvate, with L-glutamine (ThermoFisher #11965092), supplemented with 10% FBS. For quantitative assays, tissue culture dishes were treated with poly-D-lysine (ThermoFisher #A3890401).

### CRISPR/Cas9 editing

The gRNA was designed (Table S1) to cut at or one nucleotide away from 3’ of the variant site. gRNA oligonucleotides were annealed and phosphorylated in T4 Ligation buffer (NEB #B0202S) and T4 Polynucleotide Kinase (NEB #M0201). pX330 SpCas9-gRNA vector (Addgene #42230) was digested 15 mins at 37°C with BbsI and purified. The gRNA duplex was ligated into digested vector using T4 ligase (NEB #M0202) for 10 mins at room temperature and transformed into Endura electrocompetent cells (1800 V, 10 uF, 600 Ω, 1 mm cuvette, Biosearch Technologies #60242). gRNA insertion was validated by Sanger Sequencing (Azenta). Two custom single stranded oligo DNA nucleotides homology-directed repair templates (Table S1) corresponded to the desired T>G edit (one symmetric, one asymmetric). Cells were grown to 70% confluent in a 24-well culture plate and transfected using 400ng pX330 containing the gRNA, 1 uL 10uM ssODN, 1.5 uL Lipofectamine 3000, and 1 uL P3000 reagent. 48 hours after transfection, cells were plated in a 96-well plate for clonal isolation through limiting dilution and genotypes were determined via allele-specific PCR and Sanger sequencing (Azenta).

### RT-PCR and qRT-PCR

RNA was isolated (24 hours post-SSO transfection, if applicable) using the RNeasy mini kit following manufacturer’s instructions (Qiagen #74104). For end-point PCR, reverse transcription and PCR were performed as directed by SuperScriptTM III Reverse Transcriptase (Invitrogen #18080085) and EconoTaq PLUS 2x MasterMix (Biosearch Technologies #30035), and PCR product was visualized on a 1.25% agarose gel in TBE. Expression of the pseudoexon isoform or canonical isoform was quantified using probe-style qRT-PCR with the probe annealing to the *PHKG2* exon 6-pseudeoexon junction or the exon 6-exon 7 junction, respectively (Table S1). Expression was normalized to *ACTB* (Integrated DNA Technologies PrimeTime qPCR, Hs.PT.39a.22214847).

### Western Blot

Whole cell lysate was collected in 0.1% Triton X-100 (Sigma Aldrich #T8787), 150 mM NaCl, 50 mM Tris-HCl, pH 8.0, and 1X Halt Protease Inhibitor Cocktail (Thermo Scientific #78425) and quantified with the Pierce BCA Protein Assay Kit (Thermo Scientific #23227). 15 µg lysate was loaded and run in 4X Laemmli buffer (BioRad #1610747) and 10% beta-mercaptoethanol on a 4–15% Mini-PROTEAN TGX Stain-Free Protein gel (BioRad #4568083) in 1X Tris/Glycine/SDS Buffer (BioRad #1610732). Protein was transferred onto PVDF membrane for 7 minutes at 20 V on the iBlot 2 Dry Blotting System (Invitrogen #IB24002). The membrane was blocked in 4% non-fat dry milk in TBST for 1 hour at room temperature and incubated overnight with 1:2500 anti-PHKG2 Polyclonal antibody (Proteintech #15109-1-AP) or 1:2000 anti-Cyclophilin B antibody (abcam #ab16045). Membranes were washed in TBST, incubated in 1:3000 Goat Anti-Rabbit IgG H&L (HRP) in a blocking buffer for 1 hour at room temperature, and washed again in TBST before imaging using Clarity Max Western ECL Substrate (BioRad #1705062) per manufacturer instructions.

### Phosphorylase kinase enzyme activity assay

PhK enzyme activity was analyzed from frozen liver or cell lysate at the Glycogen Storage Disease Laboratory at Duke University Medical Center using standard spectrophotometric methods (44,64). Briefly, enzyme activity was measured indirectly by measuring the amount of glucose or phosphate released using Infinity glucose hexokinase (ThermoFisher #TR15421) or phosphate reagent (ThermoFisher #TR30026). Enzyme activity was expressed as μmol/min/g of liver tissue or μmol/min/mg protein from cell lysate. 2-sided *t*-tests were used to evaluate allelic differences in PhK enzyme activity.

### Doubling time investigation

300,000 cells previously grown in high glucose media (DMEM, 4.5 g/L glucose, 110 mg/L pyruvate, with L-glutamine) with 10% FBS were transferred to 6-well plates in low glucose media (DMEM with 0.5 g/L glucose, 55 mg/L pyruvate, with L-glutamine) with 10% FBS. After 48 hours, the number of cells per well was determined using an automated cell counter (Countess II FL, Thermo Fisher #AMQAF1000) and doubling time was calculated by [48 *hrs* ∗ *ln*2]/ [*ln*(*final cell count*/300,000)].

To measure rate of cell proliferation, 10,000 cells were plated in duplicate in a 96 -well plate in low glucose media. One day later, the cells were labeled with BrdU for 24 hours and BrdU was quantified by sandwich ELISA (abcam #ab126556). To measure apoptosis, 10,000 cells were plated in duplicate in a 96-well plate in high glucose media. Six hours later, they were plated in no glucose media (DMEM no glucose, with L-glutamine) supplemented with 10% FBS and added to RealTimeGlo-Annexin V detection reagent (Promega #JA1000) as directed. Luminescence and fluorescence were recorded six hours post-glucose deprivation. 2-sided *t*-tests were used to evaluate allelic differences in doubling time, BrdU incorporation, and Annexin V expression.

### Glycogen content quantification

Cells were grown in low glucose media for 48 hours, trypsinized, washed 5 times with 1x DPBS, and 600,000 cells were lysed in 0.3 N HCl. After incubating at room temperature for 5 minutes, an equal amount of Tris pH 8.0 was added. 25 uL cell lysate was plated in duplicate. The first well was treated with glucoamylase to determine glycogen content; the second well was untreated in order to assess baseline glucose content of the cells without glucoamylase treatment. 50 uL glucose detection reagent (Promega #J5051) was added to each well and mixed, and luminescence was assessed after 60 minutes incubation at room temperature. 2-sided *t*-tests were used to evaluate allelic differences in glycogen content.

### Transfection of splice-switching antisense oligonucleotides

Custom SSOs with phosphorothioate backbones and 2’ modified bases (2’-O-Methyl or 2’-O-Methoxyethyl) were obtained from Integrated DNA Technologies (Table S1). Cells were plated to 70% confluency in 24-well plates and transfected with 0-200 nM SSO or scrambled control using 2.5 uL Lipofectamine 3000 reagent and 2 uL P3000 reagent (ThermoFisher #L3000001). 24 hours post-transfection, cells were lysed for RNA extraction.

### Statistics

2-sided Student’s *t*-tests were used to compare allelic differences in gene or isoform expression, PhK enzyme activity, doubling time, glycogen, apoptosis, and BrdU incorporation. Paired 1-sided Student’s *t*-tests were used to evaluate increases in canonical *PHKG2* expression and decreases in pseudoexon expression after SSO transfection. Individual data points are taken from biologically distinct replicates rather than technical replicates. All data are presented as mean ± SEM. P-values of less than 0.05 were considered significant.

### Study approval

All studies on human subjects were approved by the Duke University Health System IRB (approval no. Pro00106476 and Pro00090878). All adult subjects and parents of minors provided written informed consent.

### Data availability

Values for all data points in graphs are reported in the Supporting Data Values file. WGS and RNA-seq data produced by this study are in the process of being submitted to dbGaP with controlled access to protect sensitive patient data.

## Author contributions

AKI, XZ, RLK, SC, MMB, JXC, MJB, WHM, ASA, GEC, PSK, and TER conceived of and designed the study. AKI, JD, RAF, and AS conducted experiments and acquired data. AKI, XZ, KP, RLK, RCR, DSB, ASA, GEC, PSK, and TER performed data analysis and interpretation. AKI, GEC, TER, and PSK wrote the manuscript, and all authors contributed to revision and editing.

## Acknowledgements

The authors gratefully acknowledge the patients and their family for their generosity in taking part in this study. We thank the following for their helpful conversations regarding experimental design, preliminary or sequencing data, interpretation of results, and/or acquisition of samples: Erin Huggins, Rebecca Gibson, Cristen Knapp, Jeong-A Lim, Schuyler Melore, Shruthi Mohan, Emi Hammond, and the Duke GSD Clinical Research team. Thank you to Azenta Life Sciences and the Duke School of Medicine for use of the Sequencing and Genomics Core Facility for performing the short- and long-read RNA-seq on patient blood and buffy coat samples. Thank you to the GTEx consortium, the ENCODE consortium, and the Thomas Gingeras lab for use of their RNA-seq data. We also thank the Duke Y.T and Alice Chen Pediatric Genetics and Genomics Research Center for its encouragement of this work. Funding was provided by NIH grants RM1HG011123, R21HG010747 and UM1HG012053.

## Notes

**<u>Conflict-of-interest statement</u>:** AKI, GEC, PSK, and TER have a patent pending (US Application no. 19/116,123).

### Competing Interest Statement

Several authors of this manuscript (AKI, GEC, PSK, TER) declare a patent pending (US Application no. 19/116,123).

