## Supplementary Figures for "Cell Modeling and Rescue of a Novel Non-coding Genetic Cause of Glycogen Storage Disease IX"

### Supplementary Material

**SpliceAI scores:** ⓘ

| Variant | Gene | ☐ = MANE Select transcript ☐ = non-coding transcript | Δ type | Δ score | position |
|---|---|---|---|---|---|
| chr16-30754626-T-G<br>intron variant<br>UCSC, gnomAD | PHKG2 ( ENSG00000156873.16 / ENST00000563588.6 / NM_000294.3)<br>protein coding MANE Select transcript (plus strand)<br>OMIM, GTEx, gnomAD, ClinGen, Ensembl, Decipher, GeneCards | | Acceptor Loss | 0.00 | |
| | | | Donor Loss | 0.02 | 131 bp |
| | | | Acceptor Gain | 0.78 | -76 bp |
| | | | Donor Gain | 0.91 | -1 bp |

**Supplementary Figure 1.** SpliceAI predicts that c.556+1069T>G creates a cryptic splice donor 1 bp downstream of the variant with high evidence and a splice acceptor 76 bp downstream of the variant with moderate evidence. This would result in an out-of-frame 76 bp pseudoexon inclusion between exons 6 and 7 of *PHKG2*.

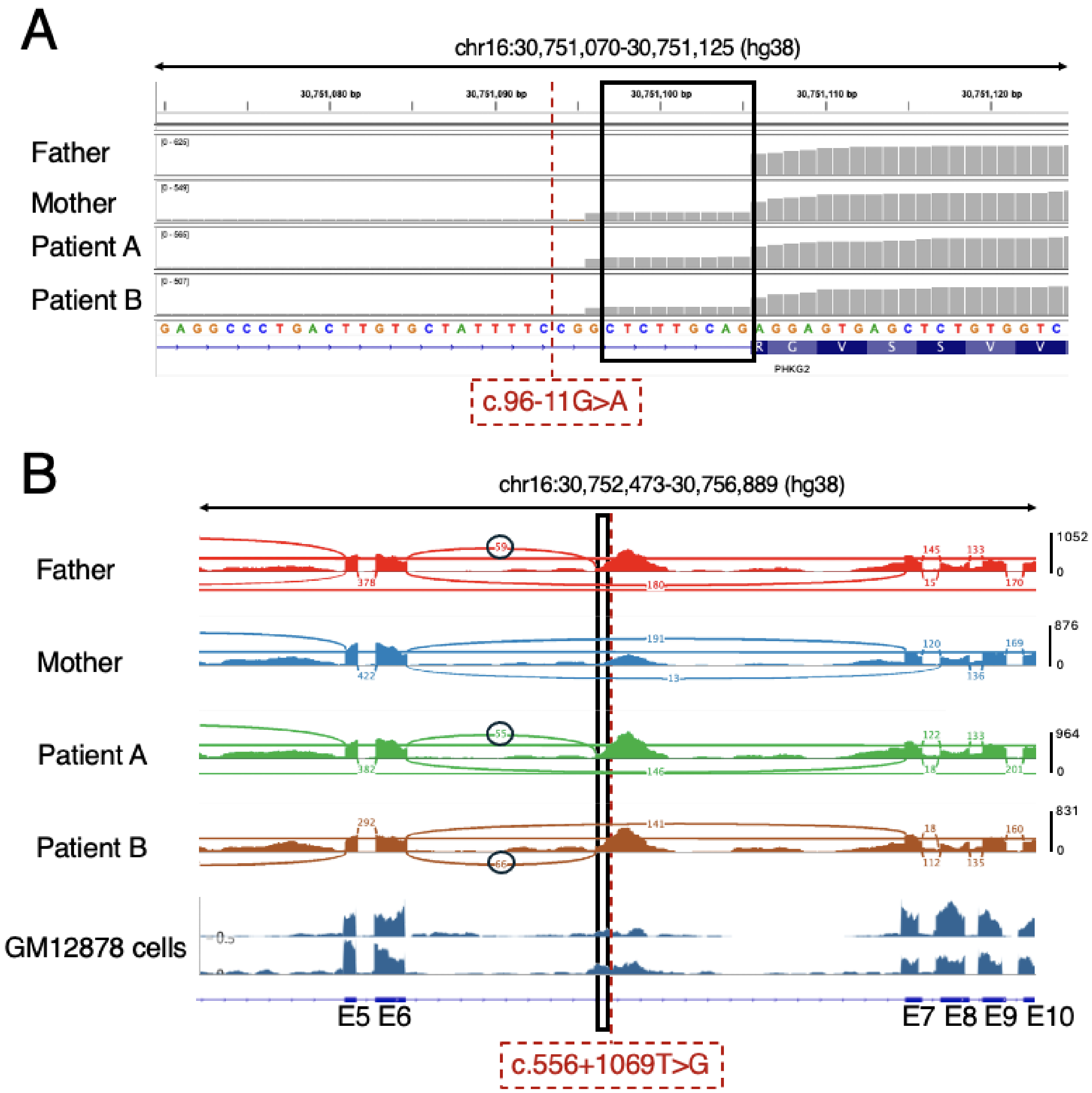


**Supplementary Figure 2.** (A) rRNA depleted RNA-seq read pileups of whole blood from Patient A, Patient B, and both parents across the exon 3 splice acceptor. This indicates that the previously known pathogenic variant c.96-11G>A causing a 9 bp exon 3 extension (boxed) carried by both siblings was inherited from the mother. (B) Sashimi plot of same individuals in (A) whole blood RNA-seq spanning the pseudoexon and surrounding exons. Exon 6-pseudoexon splice junction is found in father, Patient A, and Patient B. This indicates that the pseudoexon (boxed), and therefore c.556+1069T>G, was passed down to both siblings from the father. Read pileup peak at and downstream of the 76 bp pseudoexon is present in rRNA-depleted GM12878 human lymphoblastoid cells (ENCSR000COS) (65-67). It is not present in polyA-selected RNA-seq (Figure 2).

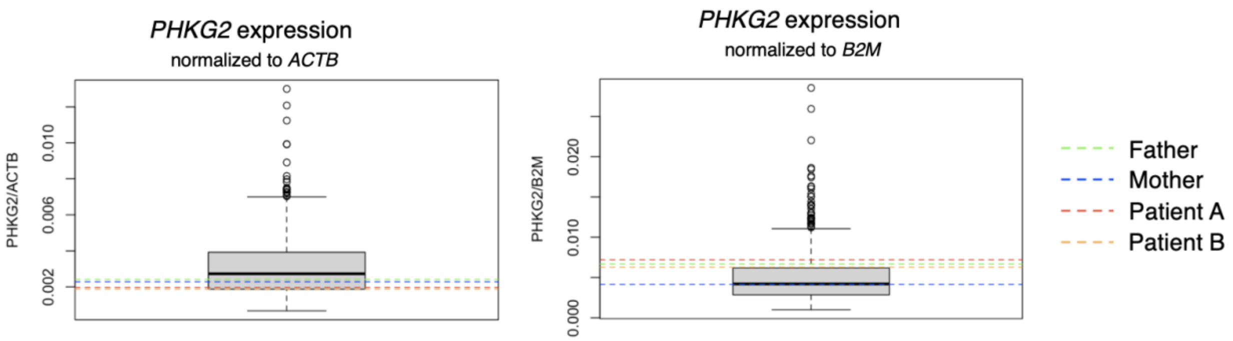


**Supplementary Figure 3.** *PHKG2* expression (TPM) normalized to housekeeping genes *ACTB* and *B2M* from whole blood RNA-seq of both siblings and parents compared to GTEx controls.

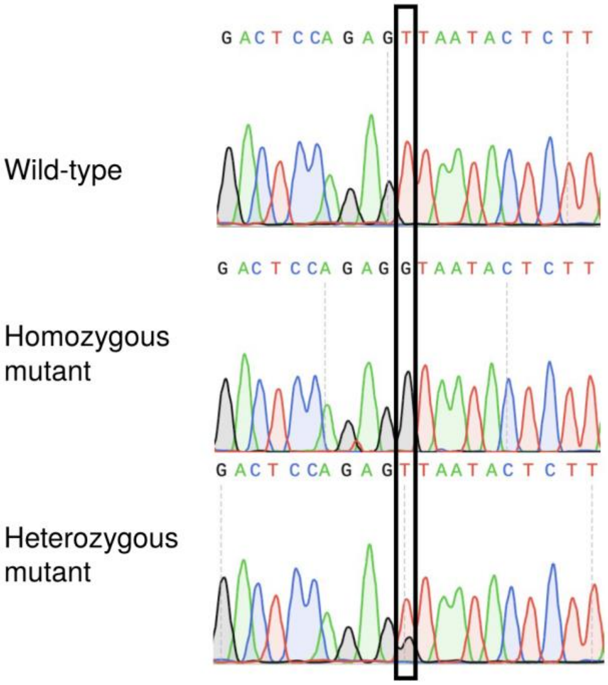


**Supplementary Figure 4.** Representative Sanger sequencing of HEK293T CRISPR-edited HEK293T clones at c.556+1069T>G (boxed).

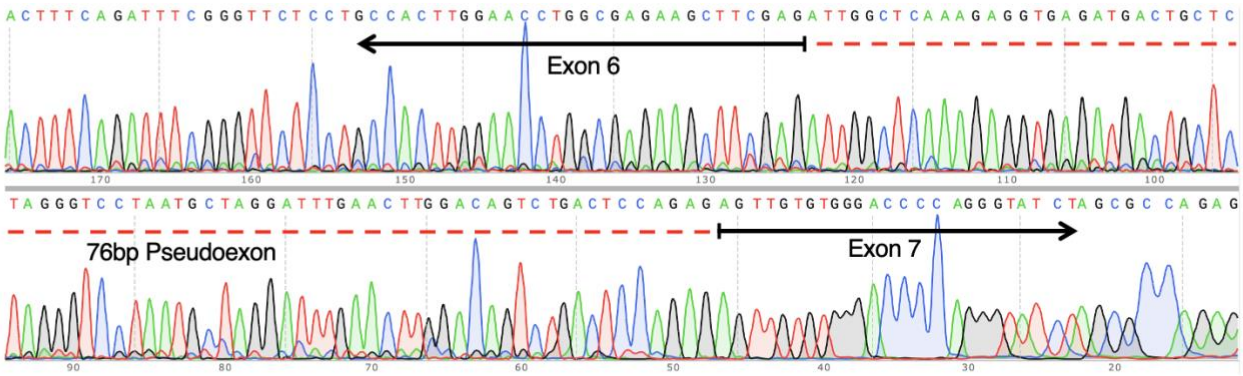


**Supplementary Figure 5**. Sanger sequencing of 380 bp RT-PCR product including PHKG2 exons 6-8 from a homozygous HEK293T edited cell clone containing c.556+1069T>G, confirming inclusion of a 76 bp pseudoexon with the predicted splice junctions. Figure shows reverse complement of sequencing data to reflect direction of transcription.

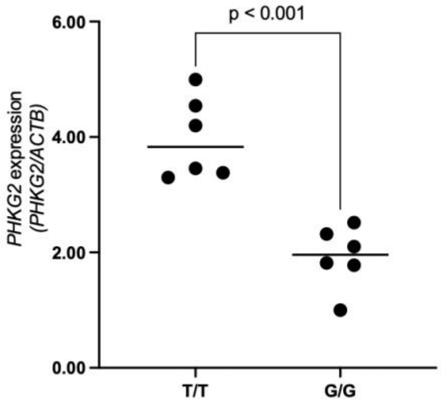


**Supplementary Figure 6.** Pseudoexon-agnostic *PHKG2* expression assay spanning exons 2-3 (ThermoFisher Taqman assay Hs04963859_m1) shows decreased expression in edited HEK293T cell lines containing the patient variant (G/G) compared to wild-type (T/T).

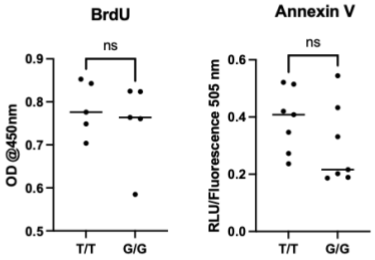


**Supplementary Figure 7.** Cell division (by BrdU) and apoptosis (Annexin V) in HEK293T cells that are wild-type (T/T allele) or containing the patient c.556+1069T>G (G/G allele).

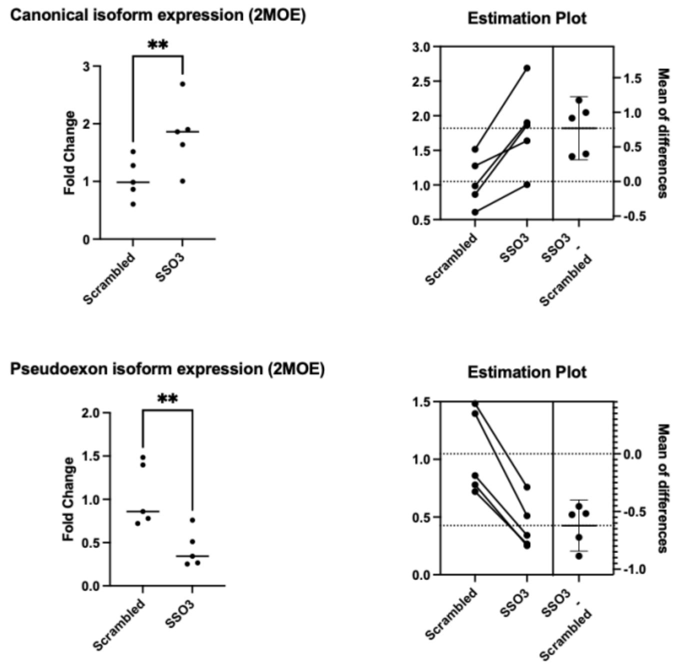


**Supplementary Figure 8.** 2'-MOE modified SSO3 and a scrambled control were transfected into 5 biologically identical GSD IX cell lines. RT-qPCR shows an increase in canonical isoform expression and a decrease in pseudoexon isoform expression in cells transfected with 2'-MOE SSO3 relative to a scrambled control 2'-MOE oligo. The estimation plots indicate that those changes are consistent across all biological replicates. (**p<0.01)
